# A geometric representation of gene-by-gene and gene-by-environment interactions on the extended complex plane

**DOI:** 10.64898/2026.06.26.734831

**Authors:** Jim Karagiannis

## Abstract

The relationship between genotypic and phenotypic variation is determined by the complex interaction of genetic and environmental factors. While statistical methods capable of detecting such interactions exist, an axiomatic mathematical framework that seamlessly describes the combined effects of genetic modifications and environmental exposures on a common scale is lacking. In this report, buffering concepts are used to construct a measurement system that enables the geometric representation of both gene-by-gene and gene-by-environment interactions on the extended complex plane (i.e., as projections on the Riemann sphere). In this manner, any such interaction, or combination thereof, can be precisely defined and quantified as the deviation from the neutral value calculated through the applicable complex transformation. When thus conceptualized, the framework’s parameterization defines the “state space” of a given measurable phenotype along both the real and imaginary dimensions, thus establishing an unambiguous and broadly applicable method for determining the phenotypic value expected upon combinatorial changes in genetic and/or environmental variables. Remarkably, by applying these methods, it is possible to quantify the effects of any gene-by-environment interaction using the equation, 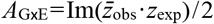, where *z*_obs_ and *z*_exp_ are complex numbers representing the observed and expected phenotypes of a given genotype expressed in terms of the buffering parameters, *ŧ* and *b*.

## Introduction

The discipline of genetics has long strived to develop the analytical tools necessary to systematically and accurately describe the relationship between genotypic and phenotypic variation[1–6]. The significance of these tools lies, not only in their potential to aid in the genetic analysis of disease phenotypes, but also in how they might be applied to better understand the mechanisms driving phenotypic diversity in populations (and thus the evolutionary history of living organisms). Three factors are thought to be key in limiting progress towards this goal: 1) the effects of interactions between genes, 2) the effects of interactions between genes and changing environments, and 3) the effects of stochastic noise. While the underlying thermodynamic basis of cellular biochemistry makes the effects of noise unavoidable, an improved methodology for the unambiguous characterization of the combinatorial effects of gene-by-gene (GxG) and gene-by-environment (GxE) interactions may be a critical step in the evolution of fully quantitative methods of genetic analysis.

A necessary prerequisite for the quantitative analysis of GxG interactions is the establishment of a set of principles that can be used to define the “neutral” phenotype (i.e., the phenotype predicted to occur in the absence of interaction). While an interaction between two genes can indeed be defined and measured as the deviation from this neutral expectation, the notion of neutral requires an *a priori* hypothesis as to how its value should be calculated. Thus, even though a variety of neutrality models have been developed, their validity, together with the circumstances under which each should be applied remains equivocal [7–9].

The analysis of GxE interactions, on the other hand, is typically based upon the definition of C. H. Waddington, who reserved the use of the term, gene-environment interaction, to “designate cases in which phenotypic effects of different genotypes are differently affected by a given environmental change” [10]. In practice, such interactions are determined by comparing the “reaction norms” of individual genotypes. A reaction norm describes the phenotype of a single genotype across a range of environments. Thus, GxE interactions can be inferred when the phenotypic response of two genotypes to the same environmental variation statistically differs [4,11–13]. Although conceptually simple, difficulties in defining and interpreting GxE interactions nonetheless exist. For example, using the previously described statistical definition, the inference of an interaction depends on whether an additive or multiplicative scale of measurement is used [13]. Thus, while the analysis of both GxG and GxE interactions have a long and rich history, ambiguities with respect to how each should be defined and/or conceptualized have not been resolved [9,12–14]. Furthermore, attempts at consolidating the analysis of GxG and GxE interactions within a common schema remain for the most part unexplored.

In this report, I present a harmonized mathematical framework capable of seamlessly describing the effects of genetic modifications and environmental exposures on a common scale. As will be described in detail below, I contend that the use of complex numbers, together with the parameterization previously developed in [15], permits the unambiguous representation of both GxG and GxE interactions on the extended complex plane (i.e., as projections on the Riemann sphere). In this way, all relevant genetic and environmental elements of the system are represented by intuitive, yet precise, geometric equivalents, thereby establishing a comprehensive and broadly applicable method for the quantitative analysis of both GxG and GxE interactions.

## Results

### Mathematical preliminaries

The methodology presented herein represents an extension of previously published work concerning the quantitative analysis of genetic buffering relationships [15]. This prior work is itself an extension of the mathematical foundation laid by B.M. Schmitt, who had earlier developed a formal and general method for the quantification of buffering action [16,17]. Before proceeding, a brief summary of the approach is necessary.

The strategy first requires the definition of three functions – the sigma function, the transfer function, and the buffer function – that together describe a measurable phenotype as it is observed in a mutant genotype relative to an arbitrarily chosen reference genotype. The sigma function, *σ* (*x*), describes the change in the quantity of interest (i.e., the measurable characteristic being observed and measured) in the chosen reference genotype (e.g., the wild-type genotype). In contrast, the transfer function, *τ*(*x*), describes the change in the quantity of interest as observed in a given mutant genotype. Finally, the buffer function, *β*(*x*), describes the change in the quantity of interest that would have been observed, if not for the effects of the mutant genotype (i.e., the portion “buffered” by the action of the mutation in question as inferred from the observation of the reference). Thus, the value of the buffer function is equal to the difference between the sigma function and the transfer function (i.e., *β*(*x*) =*σ* (*x*)-*τ* (*x*)).

Using these functions, it is possible to define four parameters, *ŧ, b, T*, and *B* (where *ŧ*=*τ*(*x*)*/σ* (*x*), *b* =*β*(*x*)*/σ* (*x*), *T=τ*(*x*)*/ β*(*x*), and *B* =*β* (*x*)*/ τ*(*x*)). Thus, the *ŧ* and *b* parameters are useful in comparing the transfer and buffer functions relative to the total change and the *T* and *B* parameters are useful in comparing the transfer and buffer functions to each other. The analysis of systems in this way is somewhat analogous to the use of probabilities (which compare a part to the whole) or odds (which compare the two parts comprising a whole) in measuring chance events.

The buffering parameters can also be used to logically derive two mutually exclusive neutrality models: one of which is applicable in cases where two alleles at different loci act “sequentially”, and another that is used when two alleles at different loci act “in-parallel” [15].

Deviations from the expected phenotype – as predicted by the respective neutrality models – are then used to define GxG interactions on a scale based on the magnitude of the deviation. Based on this framework, a GxG interaction between parallel acting alleles is defined as the case where

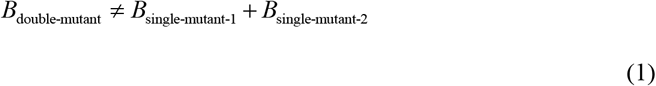

and a GxG interaction between serially acting alleles is defined as the case where

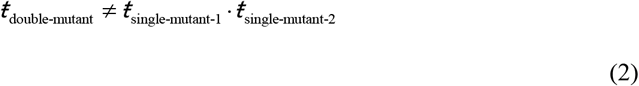

It is interesting to note that this definition provides an underlying mathematical rigor to the most common qualitative definition of gene interaction that is in current use – an interaction between alleles at different loci resulting in a phenotype distinct from that expected under the assumption of their independent action [7,18]. Thus, in short, the development of the framework described above permits the unambiguous definition and classification of gene interactions in a formal, general, and mathematical way. Further logical extensions of this framework, with the aim of seamlessly incorporating the effects of environmental variation into such analysis, form the basis of the current report and are discussed at length below.

### A geometrical description of GxG interactions on the Cartesian plane

Since the sum of *ŧ* and *b* must equal one, the set of possible phenotypes (as measured relative to that expressed by a reference genotype) can be represented geometrically by the line formed when *ŧ* and *b* are plotted on a two-dimensional Cartesian plane. Alternatively, since the product of *T* and *B* must equal one, the set of possible phenotypes can also be represented by the hyperbola formed when *T* and *B* are likewise plotted. Thus, any genotype can be mapped to the line (or hyperbola) based on its expressed phenotype. In this way, changes in phenotype that result from a genotypic change can be represented by the movement of a point along the line defined by *ŧ*+*b*=1, or by the movement of a point along the hyperbola defined by *T*·*B*=1 (**Figure 1A,B**).

**Figure 1.**
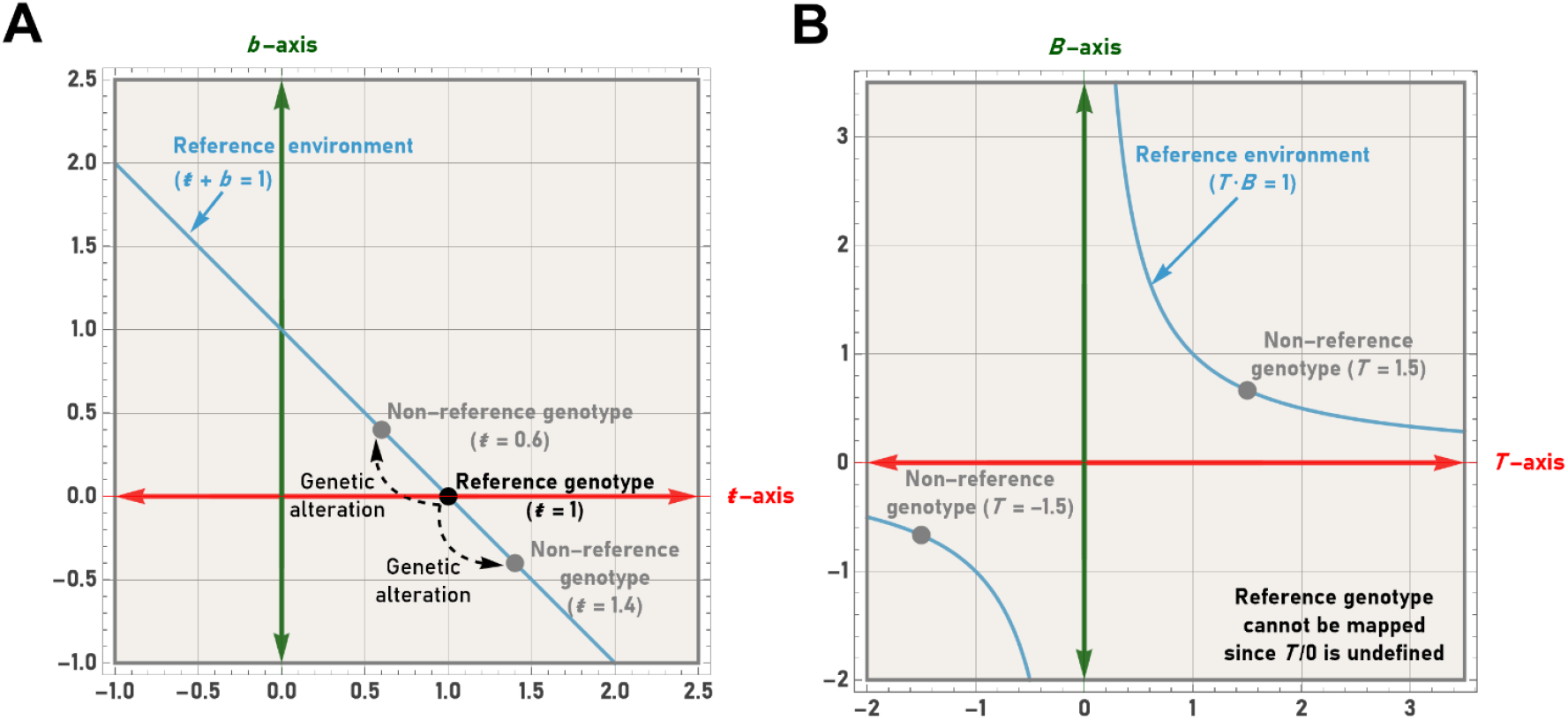
A geometric representation of the phenotypic possibilities within a given reference environment in relation to a reference genotype. When described using the transfer, buffer, and sigma functions (see text), genetic alterations can be conceptualized as the movement of a point along the line *ŧ*+*b*=1 (**A**) or along the hyperbola *T*·*B*=1 (**B**). When utilizing the buffering parameters, *ŧ* and *b*, the reference genotype can be mapped to the coordinates (1,0). When utilizing the *T* and *B* parameters, the reference genotype cannot be plotted since *T*/0 is undefined.

While these representations are capable of describing the phenotypic effects of genetic variation, there is no mechanism to incorporate the phenotypic effects of varying environmental exposures. This is because the framework implicitly assumes that the respective phenotypes expressed by a set of genotypes are measured under identical environmental conditions. Thus, the line *ŧ*+*b*=1 (or hyperbola *T*·*B*=1) represent the set of phenotypic values that are possible in a given environment (hereafter referred to as the “reference” environment). This, of course, leads to the question of how the phenotypic effects of varying environmental exposures could be represented mathematically within the framework. This extension is described in detail below.

#### Defining an environmental scaling factor

In cases where the magnitude of a phenotype is dependent upon the environment – as typically occurs, for example, when considering the growth of a unicellular microorganism at different temperatures – it is possible to define a scaling factor (*a*) that represents the fold change in the value of the sigma function in a given environment relative to an arbitrarily chosen reference environment. For example, consider an actively dividing unicellular microorganism, whose reference genotype is observed in two environments for equivalent periods of time. If the value of *σ* (*x*) in the new environment were two times that observed in the reference environment (when measured at the same instant in time), then the value of the scaling factor, *a*, would be two. Thus, *a*, can be formally defined as

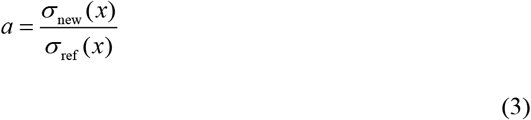

If one now assumes that the phenotypic effect of the environmental change is independent of genotype, one would expect the values of *τ* (*x*) and *β*(*x*) for any given genotype to also double in the new environment relative to the reference environment. To describe this scenario, it is first necessary to express the *ŧ* and *b* parameters as functions of both the scaling factor and the values of the respective buffering parameters in the reference environment. To avoid confusion moving forward, the values of the buffering parameters will hereafter be described where necessary using a subscript that indicates the value of the scaling factor (i.e., *ŧ*_1_ and *b*_1_ represent the values of *ŧ* and *b* in the reference environment). Using this notation the functions described above can be formally defined as:

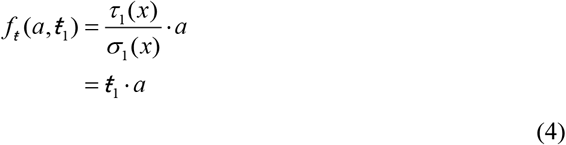

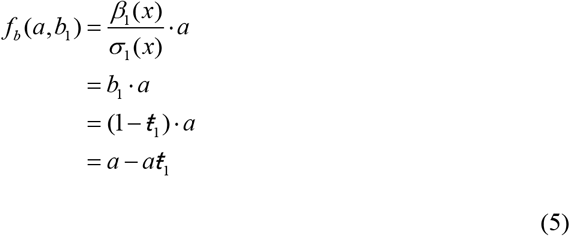

Using these definitions, it is then possible to show that

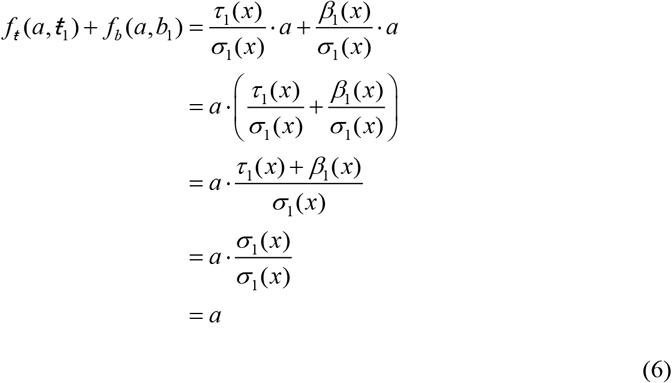

Thus, the expected set of phenotypes in environments defined by the scaling factor, *a*, can be described by the equation, *ŧ*_*a*_*+b*_*a*_ =*a*.

Similarly, it is possible to express the *T* and *B* parameters as functions of the scaling factor as follows

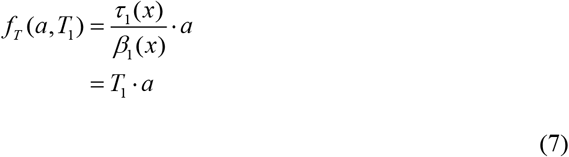

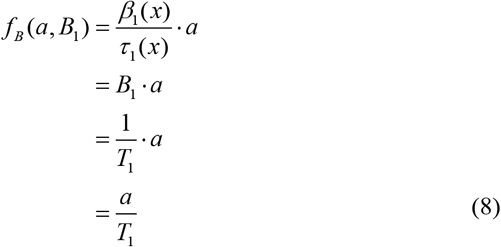

and to subsequently show that

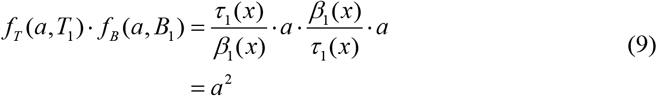

Thus, the expected set of phenotypes in environments defined by the scaling factor, *a*, can be commensurately described by the equation *T*_*a*_ ·*B*_*a*_ =*a*^*2*^. Importantly, these representations can be easily visualized using parametric plots, which formally define the respective parametric regions that describe the possible values of the relevant buffering parameters for any value of *a* (**Figure 2A,B**).

**Figure 2.**
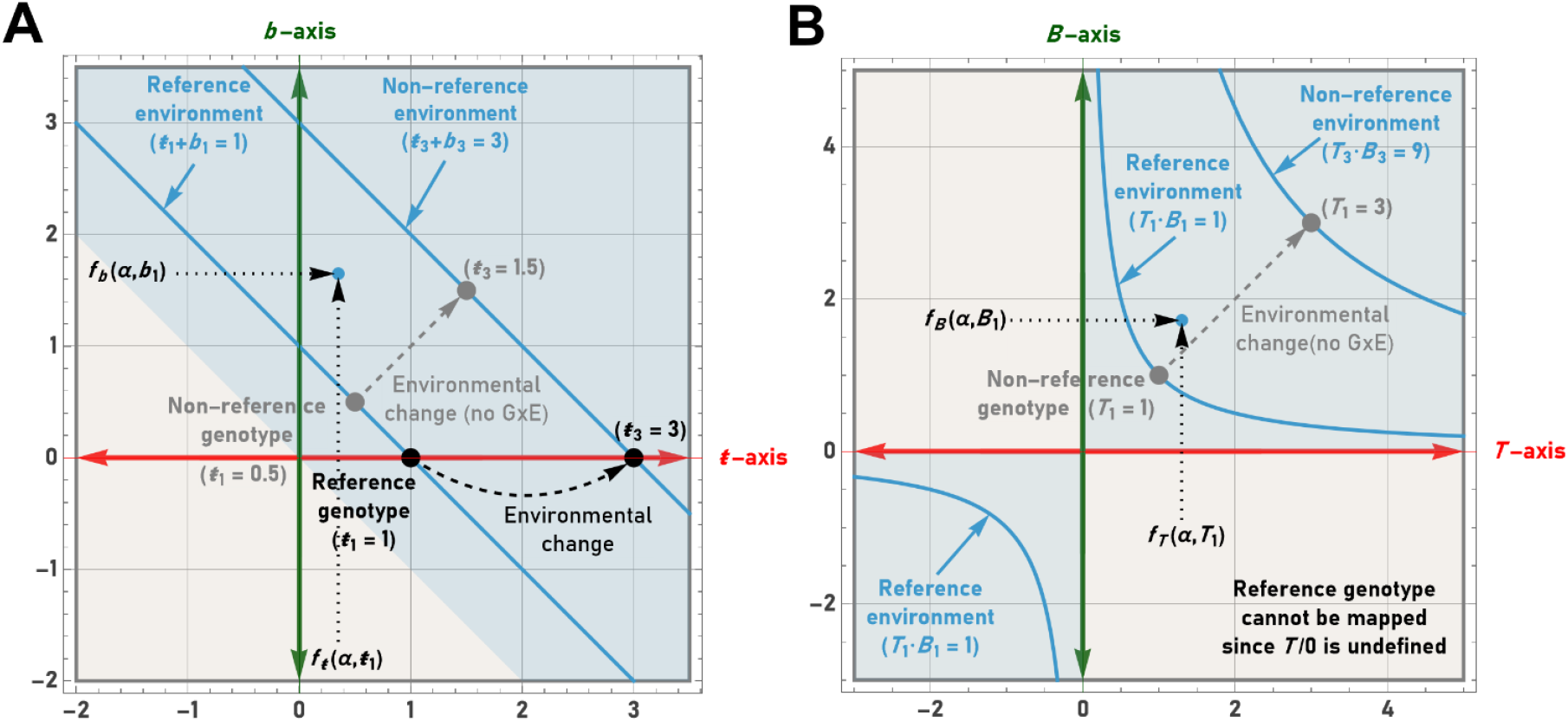
The representation of environmental changes as the scaling of the reference line or reference hyperbola. When described as functions of the environmental scaling factor, *a*, it is possible to utilize the buffering parameters to describe the phenotypic possibilities (blue shading) within a given environment using the equations, *ŧ*_*a*_*+b*_*a*_ =*a* (**A**) or *T*_*a*_ ·*B*_*a*_ =*a*^*2*^ (**B**). The reference genotype can be mapped to the coordinates (*a*, 0) when using the *ŧ* and *b* parameters, but cannot be plotted when using the *T* and *B* parameters since division by zero is undefined. Environmental changes can thus be described as the scaling of the reference line or reference hyperbola (as determined by the value of *a*).

In terms of the hypothetical unicellular microorganism described earlier, the set of possible phenotypes expressed by genetic variants in the new environment (relative to the reference genotype) can now be represented by the line defined by *ŧ*_2_ +*b*_2_ =2 or by the hyperbola defined by *T*_2_ ·*B*_2_ =4. Similarly, if the value of *σ* (*x*) decreased two-fold upon observing a reference genotype in a given environment vs. a reference environment, then the expected set of phenotypes in this environment could be represented by the line defined by *ŧ*_0.5_ +*b*_0.5_ =0.5 or by the hyperbola defined by *T*_0.5_ ·*B*_0.5_ =0.25. It should also be noted that scenarios in which *a*=0 simply reflect the fact that no phenotype is observable (i.e., the organism under observation has reached equilibrium with its environment and can no longer be differentiated from its surroundings). The equivalencies of the buffering parameters with scaling are provided in **Table 1** for reference.

**Table 1:** Buffering parameter equivalencies with scaling.

| **$t_a$ equates to:** |  | **$b_a$ equates to:** |  | **$T_a$ equates to:** |  | **$B_a$ equates to:** |  |
| --- | --- | --- | --- | --- | --- | --- | --- |
| $t_a$ | $t_1 \cdot a$ | $a - t_a$ | $a - a \cdot t_1$ | $\frac{a \cdot t_a}{a - t_a}$ | $\frac{a^2 \cdot t_1}{a - a \cdot t_1}$ | $\frac{a^2 - a \cdot t_a}{t_a}$ | $\frac{a^2 - a^2 \cdot t_1}{a \cdot t_1}$ |
| $a - b_a$ | $a - a \cdot b_1$ | $b_a$ | $b_1 \cdot a$ | $\frac{a^2 - a \cdot b_a}{b_a}$ | $\frac{a^2 - a^2 \cdot b_1}{a \cdot b_1}$ | $\frac{a \cdot b_a}{a - b_a}$ | $\frac{a^2 \cdot b_1}{a - a \cdot b_1}$ |
| $\frac{a \cdot T_a}{a + T_a}$ | $\frac{a^2 \cdot T_1}{a + a \cdot T_1}$ | $\frac{a}{1 + \frac{T_a}{a}}$ | $\frac{a}{1 + T_1}$ | $T_a$ | $T_1 \cdot a$ | $\frac{a^2}{T_a}$ | $\frac{a}{T_1}$ |
| $\frac{a}{1 + \frac{B_a}{a}}$ | $\frac{a}{1 + B_1}$ | $\frac{a \cdot B_a}{a + B_a}$ | $\frac{a^2 \cdot B_1}{a + a \cdot B_1}$ | $\frac{a^2}{B_a}$ | $\frac{a}{B_1}$ | $B_a$ | $B_1 \cdot a$ |

### A mathematical representation of the buffering parameters using complex numbers

While possible to view and manipulate the elements of the current framework using only real numbers (ℝ), they are more easily dealt with mathematically when conceptualized as complex numbers (C) on the Argand plane. Essentially, since complex numbers form a field (in the mathematical sense of the term) they provide a powerful means by which to represent rotations and scaling within two-dimensional space, thereby simplifying calculations and allowing the use of well-developed methods for their visualization and analysis [19–22]. As will be shown in subsequent sections, these tools become necessary for further developing the framework so that it is able to encompass the characterization of GxE interactions. Thus, in the same way that complex numbers are used to aid in the analysis of systems in physics and engineering, they too can be used to aid in the analysis of genetic systems. However, before these extensions can be properly explored, it is necessary for the framework to be conceptualized using the complex numbers.

#### Conceptualizing buffering parameter values as complex numbers on the Argand plane

To conceptualize the current framework in C it is first necessary to define a given phenotype as an individual complex number. This can be achieved by letting *ŧ* and *b* represent the real and imaginary parts of *z* (i.e., by letting *ŧ*=Re(*z*) and *b*=Im(*z*)). In this way, the phenotype expressed by any genotype can be described in rectangular format as follows

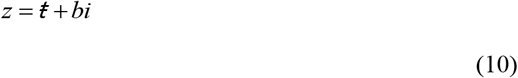

It is then possible to define z_*a*_ as

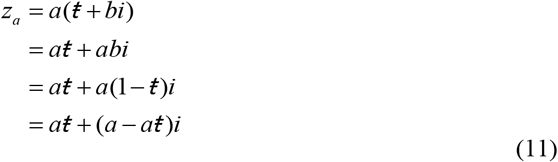

Given the definition of *z*_*a*_, it is then possible to state that

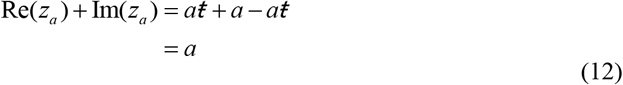

thereby, recapitulating Eq. (6).

Similarly, one could let *T* and *B* represent the real and imaginary parts of *Z*. In this way, the phenotype expressed by any genotype can be described in rectangular format as follows

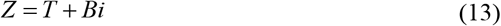

It is then possible to define *Z*_*a*_ as

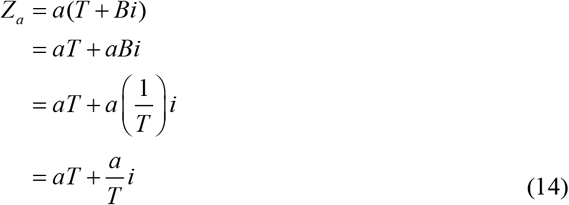

Given the definition of *Z*_*a*_, it is then possible to state that

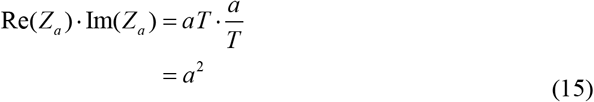

thereby recapitulating Eq. (9).

Thus, when conceptualized in C, the set of phenotypes expected in any given environment (assuming no GxE interaction) can be defined by the set of complex numbers satisfying the equations below:

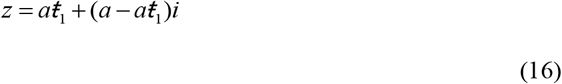

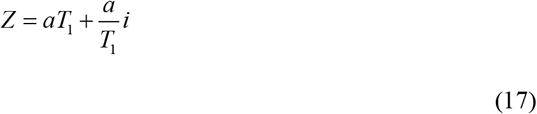

where *z* or *Z* denotes the complex number defining a given phenotypic value, *i* denotes the imaginary unit, *a* denotes the environmental scaling factor, and *ŧ*_*1*_ and *T*_1_ represent the respective values of the *ŧ* and *T* parameters in the reference environment.

Using this framework, any phenotypic value exhibited by any genotype in any environment (in relation to a given reference genotype and a given reference environment) can be represented by a complex number on the Argand plane (**Figure 3A,B**). In other words, the current framework defines the “state space” of a given phenotype (i.e., the space of all possible phenotypes along <u>both</u> the real and imaginary dimensions). In this way, information regarding the transfer and non-transfer (i.e., buffering) of the the quantity of interest, *x*, is provided in relation to both genetic and environmental factors.

**Figure 3.**
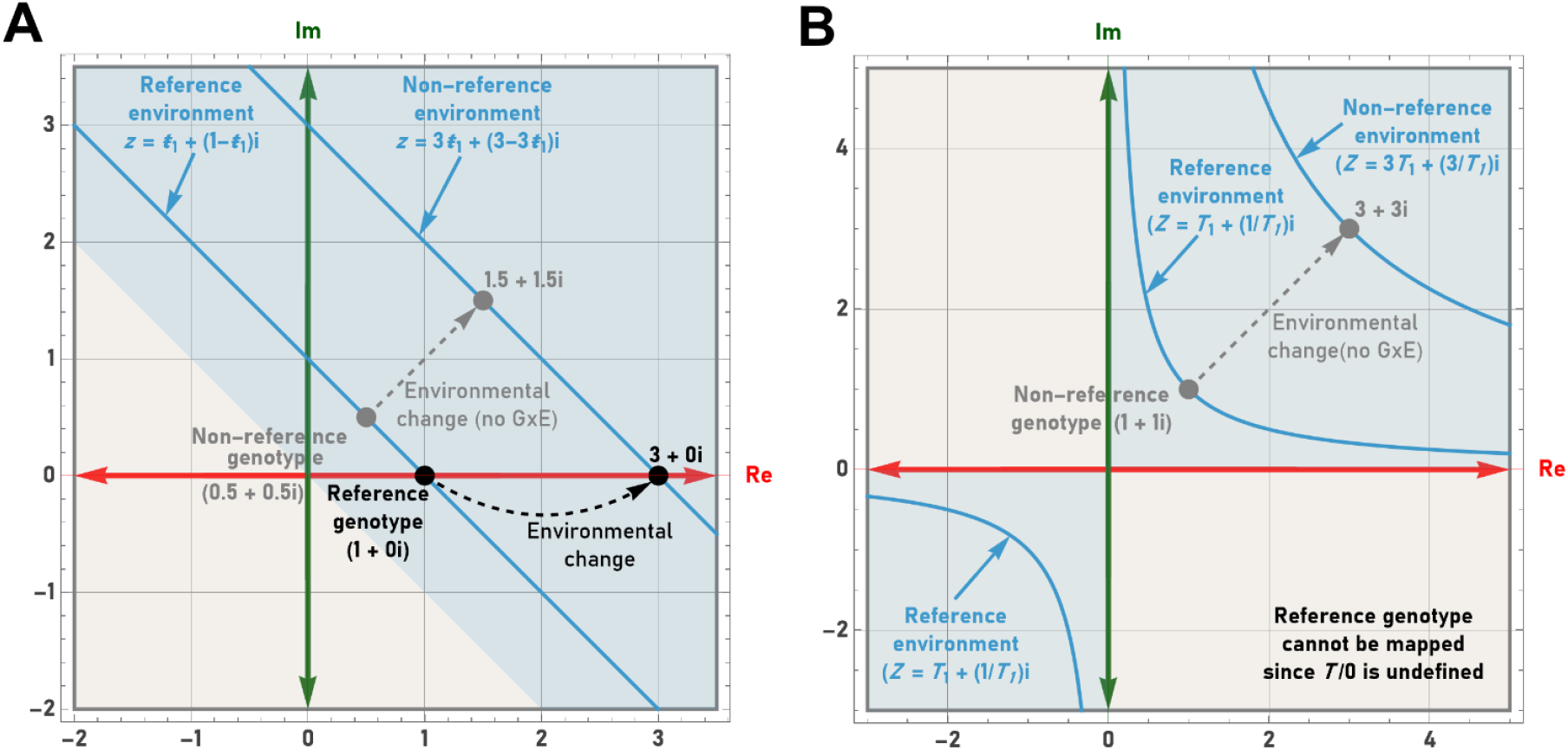
Representing genetic modifications and environmental exposures using complex numbers plotted on the Argand plane. (**A**) By letting *ŧ* and *b* represent the real and imaginary parts of *z* it is possible to define the phenotypic possibilities within a given environment (blue shading) by the set of complex numbers satisfying the equation, *z*= *aŧ*_1_+(*a*-*aŧ*_1_)*i*. The location of the reference genotype on the Argand plane in this case is determined by the complex number *z*=*a* +0*i*. (**B**) By letting *T* and *B* represent the real and imaginary parts of *Z* it is possible to define the phenotypic possibilities within a given environment by the set of complex numbers satisfying the equation, *Z*=*a*/*T* _1_ +(*a*/*T*_1_)*i*. The reference genotype cannot be mapped when using the *T* and *B* parameters since division by zero is undefined.

### A formal and general definition of GxE interaction

As described above, the current framework is capable of representing the effects of genetic and environmental changes in the absence of GxE interactions. To extend the framework so that it is able to encompass cases that do involve GxE interactions, one must first recognize the fact that complex numbers possess both a magnitude (also known as the modulus) and a phase (also known as the argument). The magnitude is defined as the absolute value of the given complex number, *z*, whereas the phase is defined as the angle (measured counterclockwise) from the positive real axis to the line connecting the origin to *z* on the Argand plane. Since GxE interactions, by definition, refer to cases in which the phenotypic effects of different genotypes are differentially affected by the same environmental change (or vice-versa), it is possible to define and quantify the effects of a GxE interaction using the phase difference in the expected value of *z* (*z*_exp_) and the observed value of *z* (*z*_obs_) in that environment.

For example, consider a reference environment (defined by the line Re(*z*) +Im(*z*)=1) where the reference genotype (WT) is mapped by definition to

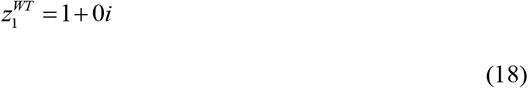

In the new environment (defined by the line Re(*z*)+Im(*z*)=*a*), the reference genotype is mapped, again by definition, to

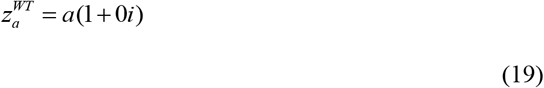

In the absence of interaction, the mutant genotype (MT) would thus be expected to map to

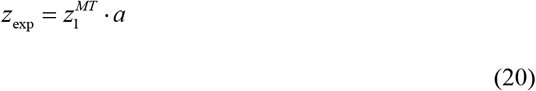

Since multiplying a complex number, *z*, by a scalar alters only the magnitude of *z*, the phase of *z*_exp_ remains equal to *z*_1_.

If, on the other had, the transfer of the quantity of interest is less than expected (i.e., the interaction between the genotype and the environment *interferes* with the transfer), or if the transfer of the quantity of interest is greater than expected (i.e., the interaction between the genotype and the environment *facilitates* the transfer), then the phase of *z*_obs_ and *z*_exp_ would no longer be equal (thereby revealing the existence of a GxE interaction). To reiterate, in cases where the transfer is indeed greater or lesser than expected, it must be the case that the corresponding difference flowed (in an abstract sense) into or out of the buffer compartment. Geometrically, this deviation from the expected outcome can be viewed as the movement of the point defined by *z*_exp_ to the point along the line Re(*z*)+Im(*z*)=*a* that accounts for the difference, thus defining the point *z*_obs_ (which must therefore possess a phase distinct from *z*_exp_) (**Figure 4A**).

**Figure 4.**
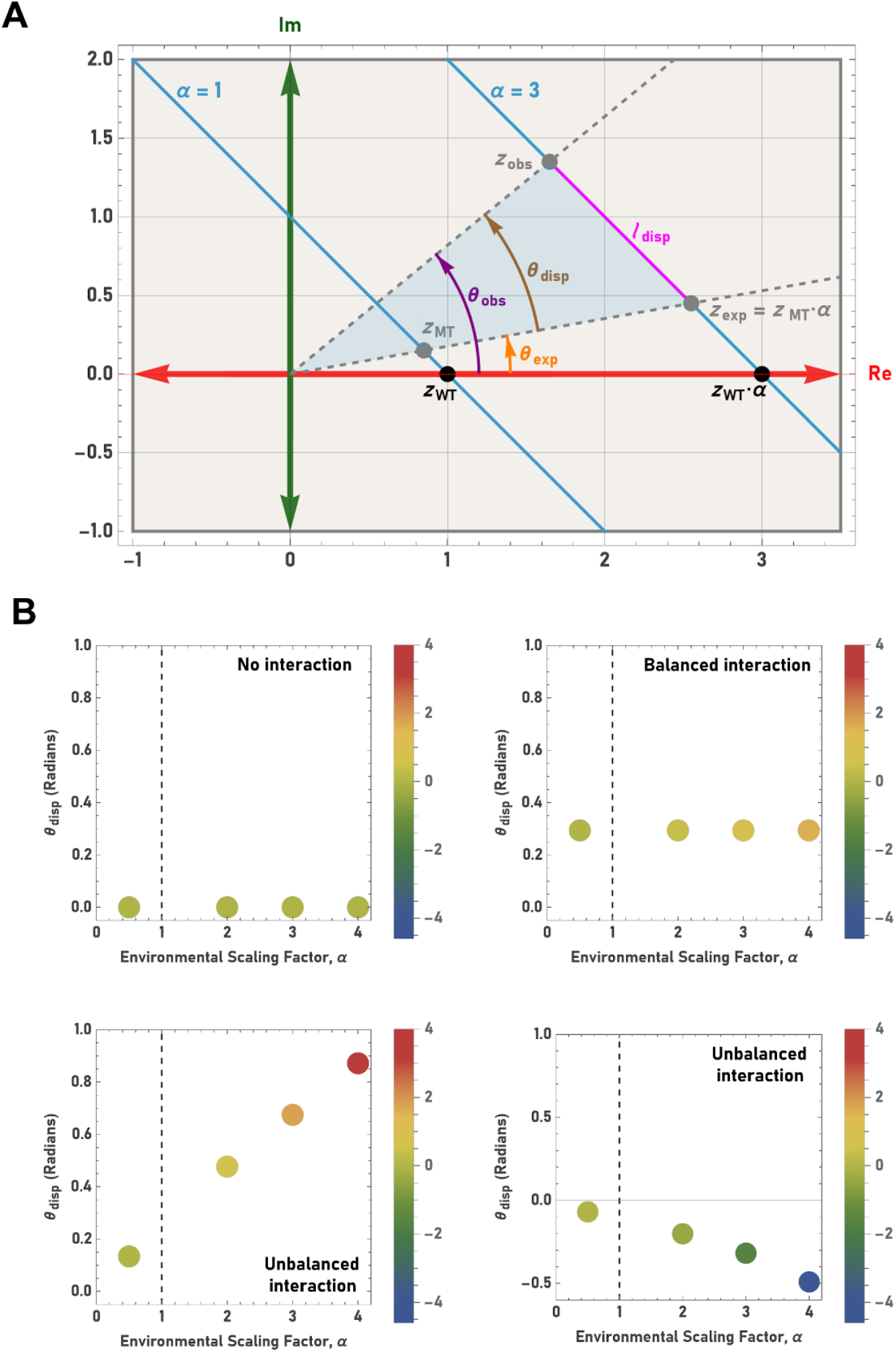
A formal and general definition of GxE interaction using the buffering parameters, *ŧ* and *b*. (**A**) It is possible to unambiguously quantify the effects of GxE interactions by taking advantage of the magnitude and phase differences in the observed value of *z* (*z*_obs_) and the expected value of *z* (*z*_exp_) on the Argand plane. The value of *z*_exp_ is equal to the product of the scaling factor, *a*, and the value of *z* that describes the mutant phenotype in the reference environment (i.e., *z*_exp_=*z*_MT_·*a*). In so doing, it is possible to define i) the linear displacement (*ℓ*_disp_) between *z*_obs_ and *z*_exp_, ii) the angular displacement between *z*_obs_ and *z*_exp_ (*θ*_disp_), and iii) the area of the triangle formed by the points representing the origin, *z*_obs_, and *z*_exp_ (blue shading). The calculated area of the triangle (see text) amalgamates the effects of both environmental changes (i.e., changes in *a*) and genetic changes (i.e., deviations in phase between *z*_obs_ and *z*_exp_). (**B**) The metrics described in (A) can be integrated into a single plot to provide a visual representation of the effects of GxE interactions. The value of the scaling factor, *a*, is shown on the x-axis, the value of *θ*_disp_ is shown on the y-axis, and the value of the area (*A*_GxE_) is represented by a color gradient. The nature of any GxE interaction can thus be described in terms of both environmental and genetic factors. The absence of interaction is characterized by a null phase difference and area. Balanced interactions maintain the same angular displacement across environments, indicating the genetic effect remains constant. Unbalanced interactions are characterized by changes in the angular displacement, indicating that the environmental change either facilitates or interferes with the genetic effect (or vice-versa).

Accordingly, it is possible to quantify the magnitude of the deviation by defining a complex transformation that takes *z*_exp_ as input and outputs the value of *z*_obs_. One way in which to achieve this is to identify a complex number (denoted *p*_GxE_) that when added to *z*_exp_ gives *z*_obs_. In this case, the complex transformation (denoted *I*_Tr_) is a translation, and can be formally defined as

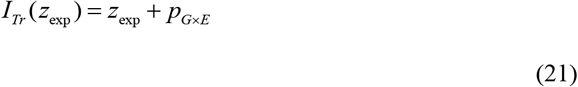

where

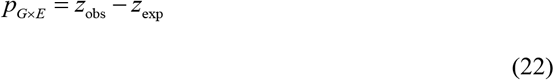

A second way in which to achieve the same goal is to identify a complex number (denoted *q*_GxE_) that when multiplied by *z*_exp_ gives *z*_obs_. In this case the transformation (*I*_RD_) combines both a rotation and a dilation, and is defined by

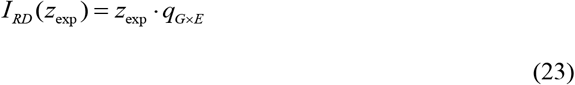

where *q*_GxE_ can be defined (in exponential form) as

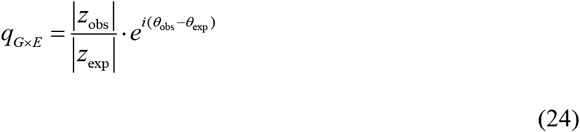

In either case, the values of *p*_GxE_ or *q*_GxE_ can be used to unambiguously define the deviation between *z*_obs_ and *z*_exp_. Similarly, when using the *T* and *B* parameters, it is possible to define the analagous complex numbers as follows:

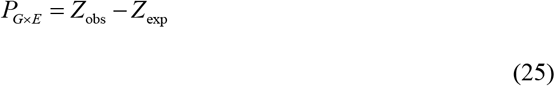

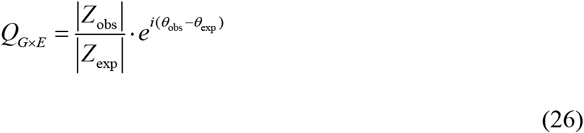

While either *p*_GxE_, *q*_GxE_, *P*_GxE_, or *Q*_GxE_ can be used to precisely define the effects of a given GxE interaction, the obtained values are somewhat nonintuitive and difficult to conceptualize, especially when attempting to compare numerous instances of GxE interactions in varying environments. For this reason, I define alternate measures below that more intuitively capture the magnitude of the deviation between *z*_obs_ and *z*_exp_ (or between *Z*_obs_ and *Z*_exp_), and that can be used to generate simple graphical representations of GxE interactions.

### Quantifying the deviation between *z*_obs_ and *z*_exp_

One simple way in which to quantify the deviation between *z*_obs_ and *z*_exp_ is to determine the linear displacement (*ℓ*_disp_) between the points on the Argand plane. This displacement is defined by

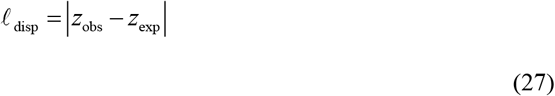

Geometrically, this measure represents the length of the line joining the points *z*_obs_ and *z*_exp_ (**Figure 4A**).

The second alternative quantifies the angular displacement (*θ*_disp_) that exists between *z*_obs_ and *z*_exp_. This displacement is defined by

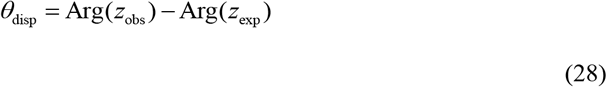

Geometrically, this measure represents the angle formed by the line joining the origin to *z*_obs_, and the line joining the origin to *z*_exp_ (**Figure 4A**).

Importantly, it is possible to use the Cosine Law (a trigonometric identity) to calculate the value of *ℓ*_disp_ given the value of *θ*_disp_, or to calculate the value of *θ*_disp_ given the value of *ℓ*_disp_. In fact, since the Cosine Law acts a bijective mapping, it is possible to conclude that there is a one-to-one correspondence between the values of *ℓ*_disp_ and *θ*_disp_. Since the Cosine Law relates the side lengths of a triangle (in this case the triangle formed by joining the points defined by the origin, *z*_obs_, and *z*_exp_) to the cosine of one of the angles (in this case *θ*_disp_), it is possible to define the linear displacement as

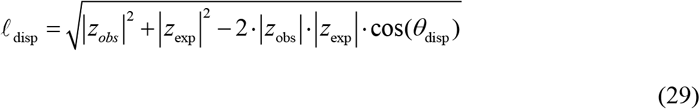

Similarly, the angular displacement can be defined as

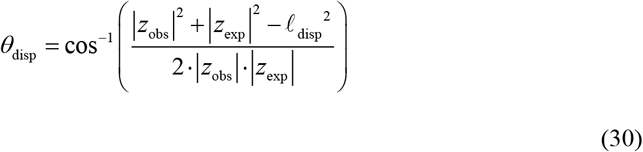

Furthermore, it is also possible to amalgamate the information provided by *ℓ*_disp_ and *θ*_disp_ by defining the area (*A*_GxE_) of the triangle formed by the origin, *z*_obs_, and *z*_exp_. This can be achieved in terms of *θ*_disp_ using the side-angle-side method

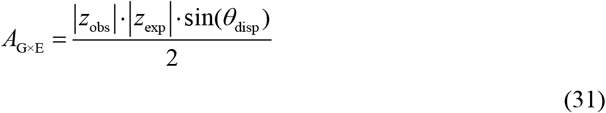

or in terms of *ℓ*_disp_ by using Heron’s Law, which first requires the calculation of the semi-perimeter (*s*)

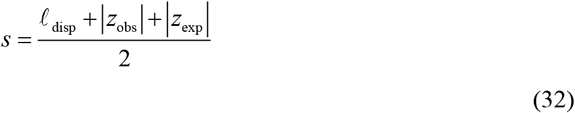

that is used, in turn, to calculate the area (*A*_GxE_)

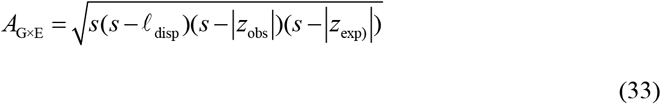

Alternatively, by taking advantage of the principles of complex geometry, it is possible to determine the area by simply applying the method of Zwicker [23] as follows

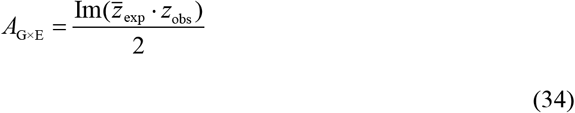

Remarkably, the GxE effect can thus be unambiguously determined through a simple calculation involving only the complex numbers representing the observed and expected phenotypes of a given genotype (expressed in terms of the buffering parameters, *ŧ* and *b*).

Ultimately, the metrics described above can be integrated within a single plot to provide an unambiguous visual representation of the effects of any GxE interaction. In such plots, the value of the scaling factor, *a*, is represented by the *x*-coordinate, the value of *θ*_disp_ or *ℓ*_disp_ is represented by the *y* coordinate, and finally, the value of *A*_GxE_ is represented by a color gradient (**Figure 4B**). In so doing, it is possible to precisely define the nature of any GxE interaction in terms of the effects of both environmental and genetic factors.

### Quantifying the deviation between *Z*_obs_ and *Z*_exp_

The deviation between *Z*_obs_ and *Z*_exp_ can be determined using a strategy analagous to that used in the preceding section. However, when dealing with *Z*_obs_ and *Z*_exp_, the hyperbolic nature of the curve defined by the *T* and *B* parameters must be taken into consideration (**Figure 5A**). While this fact complicates the calculation of the distance and area metrics, the calculation of the angular displacement (*Θ*_disp_) can still be simply defined as

**Figure 5.**
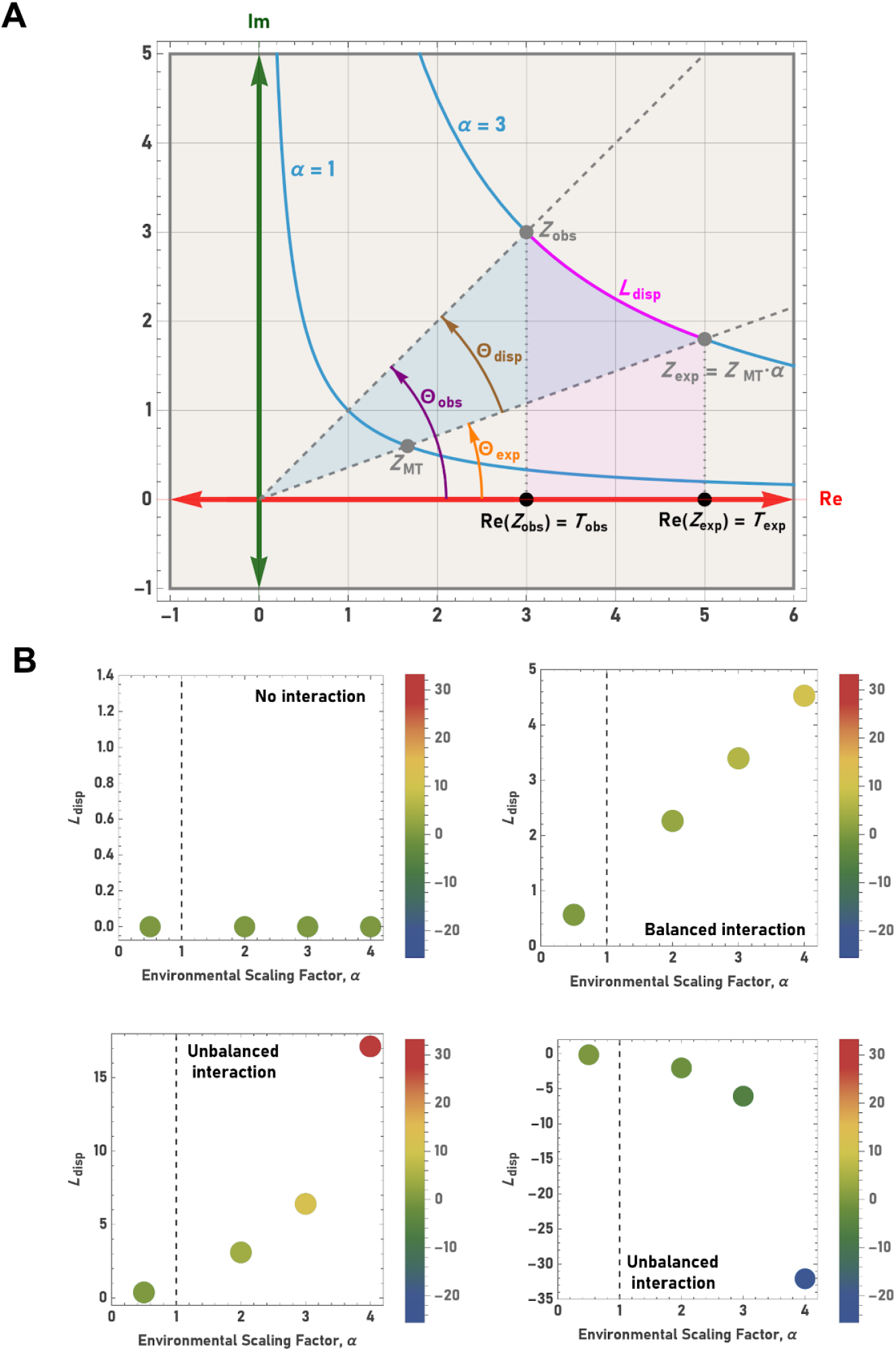
A formal and general definition of GxE interaction using the buffering parameters, *T* and *B*. (**A**) It is possible to unambiguously quantify the effects of GxE interactions by taking advantage of the magnitude and phase differences in the observed value of *Z* (*Z*_obs_) and the expected value of *Z* (*Z*_exp_) on the Argand plane. The value of *Z*_exp_ is equal to the product of the scaling factor, *a*, and the value of Z that describes the mutant phenotype in the reference environment (i.e., *Z*_exp_=*Z*_MT_·*a*). In so doing, it is possible to define i) the linear displacement (*L*_disp_) between *Z*_obs_ and *Z*_exp_ (i.e., the arc length between *Z*_obs_ and *Z*_exp_), ii) the angular displacement between *Z*_obs_ and *Z*_exp_ (*Θ*_disp_), and iii) the area of the hyerbolic sector formed by the points representing the origin, *Z*_obs_, and *Z*_exp_ (blue shading). Importantly, the area of this hyerbolic sector is equivalent to the area under the hyperbolic curve between *T*_obs_ and *T*_exp_ (pink shading). This calculated area (see text) amalgamates the effects of both environmental changes (i.e., changes in *a*) and genetic changes (i.e., deviations in phase between *Z*_obs_ and *Z*_exp_). (**B**) The metrics described in (A) can be integrated into a single plot to provide a visual representation of the effects of GxE interactions. The value of the scaling factor, *a*, is shown on the x-axis, the value of *L*_disp_ is shown on the y-axis, and the value of the area (*Λ*_GxE_) is represented by a color gradient. The nature of any GxE interaction can thus be described in terms of both environmental and genetic factors. The absence of interaction is characterized by a null phase difference and area. Balanced interactions exhibit a linear relationship between *L*_disp_ and area, indicating the genetic effect remains constant. Unbalanced interactions are characterized by non-linear relationships between *L*_disp_ and area, indicating that the environmental change either facilitates or interferes with the genetic effect (or vice-versa).

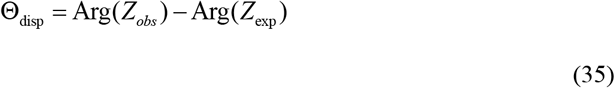

Thus, *Θ*_disp_ represents the angle formed by the line joining the origin to *Z*_obs_, and the line joining the origin to *Z*_exp_ (**Figure 5A**).

Regarding the derivation of an analogous distance metric, the hyperbolic nature of the curve necessitates a modification to the logic that must be employed. This is because, unlike *ℓ*_disp_, which calculates the straight line distance between *z*_obs_ and z_exp_, the distance between *Z*_obs_ and *Z*_exp_ must be defined as the length of the arc of the hyperbola between *Z*_obs_ and *Z*_exp_ (**Figure 5A**). As shown below, this can be accomplished using established geometrical principles together with standard integral calculus (since it is only necessary to evaluate the integral of the real part with respect to the real axis):

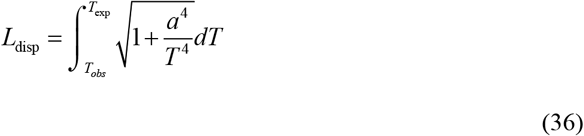

Unfortunately, this integral is non-elementary and must be solved using numerical methods (i.e., it cannot be expressed as a finite combination of basic functions).

Regarding the derivation of an analogous area metric, it is again possible to use standard integral calculus. This is because the area of the hyerbolic sector defined by joining the origin and the points defined by *Z*_obs_ and *Z*_*e*xp_ is equivalent to the area under the hyperbolic curve between *T*_obs_ and *T*_exp_ (**Figure 5A**). For this reason, it is possible to calculate the GxE effect as follows:

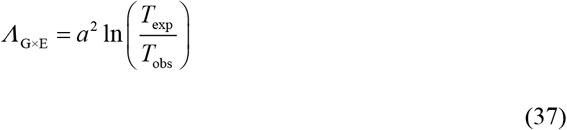

In this way, the GxE effect can again be unambiguously determined through a simple calculation; in this case, one involving only the value of the scaling factor, together with the values of *T*_exp_ and *T*_obs_.

In the same manner in which the analagous metrics were integrated into a single plot in the preceding section, these metrics can also be integrated into a single plot to provide an unambiguous visual representation of the effects of any GxE interaction. In this case, the value of the scaling factor, *a*, is represented by the *x*-coordinate, the value of *Θ*_disp_ or *L*_disp_ is represented by the *y*-coordinate, and finally, the value of *Λ*_GxE_ is represented by a color gradient (**Figure 5B**). Again, these plots can be used to precisely define the nature of any GxE interaction in terms of the effects of both environmental and genetic factors.

### Projections of the Argand plane onto the Riemann sphere

The introduction of discontinuities is a difficulty that is encountered when considering scenarios involving perfect transfer or perfect buffering. This is due to the fact that the *T* and *B* parameters, respectively, become undefined in such cases as a result of division by zero. Moreover, scenarios involving perfect buffering or perfect transfer cannot be plotted since the real and imaginary axes of the Argand plane are asymptotes of the hyperbola, Re(*Z*)·Im(*Z)*=*a*^2^. Fortunately, this limitation can be overcome – just as it is when this circumstance arises in the physical or engineering sciences – by using the extended complex plane, C_∞_ (where division by zero is indeed defined).

The extended complex plane is formed by the union of the complex plane, C, and a single point at infinity (i.e., C_∞_ = C ∪ {∞}). This can be visualized by imagining the two-dimensional complex plane being wrapped around a sphere (referred to as the Riemann sphere) whose south pole is positioned at the complex origin. The mapping of points from the complex plane to the surface of the sphere is peformed using a stereographic projection. Geometrically, this projection, *P*(*z*), can be visualized as the point of intersection between the surface of the sphere and a line drawn from the north pole to the point on the complex plane, *z*, that is being projected. In this way, every point on the complex plane can be mapped to a single, unique point on the surface of the sphere.

When using the *ŧ* and *b* parameters, the coordinates used to describe the location of the projected point on the sphere consist of the *ŧ*-axis coordinate, the *b*-axis coordinate, and the coordinate of a third axis (the *w*-axis), which defines the third spatial dimension that is needed to incorporate the sphere (and which is perpendicular to both the *ŧ*-axis and *b*-axis). Similarly, when using the *T* and *B* parameters, the coordinates used to describe the location of the projected point consist of the *T*-axis coordinate, the *B*-axis coordinate, and the coordinate of a third axis (the *W*-axis) that is perpendicular to both the *T*-axis and *B*-axis. The coordinates of the projection of a complex number, *z*, for a Riemann sphere whose south pole is located at (0,0,0) and whose north pole is located at (0,0,1) are defined as follows

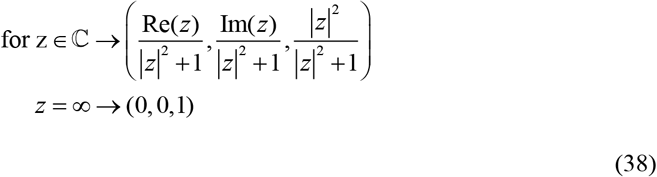

A consequence of this method of stereographic projection is that as the magnitude of a complex number approaches infinity (i.e., as |*z*| → ∞) its projection approaches the north pole, irrespective of its phase. In other words, all “paths” extending outwards from the complex origin, in any direction, converge at a single point defined by the north pole. Thus, the extended complex plane forms a closed and bounded space (formally referred to as a “compactified” space) where the north pole corresponds to infinity and the south pole corresponds to the complex origin (**Figures 6A,B**). Therefore, when considered in the context of the *T* and *B* parameters, this conceptualization in C_∞_ allows cases of perfect transfer or perfect buffering to be mapped to infinity instead of being undefined (as they would be if conceptualized in C).

**Figure 6.**
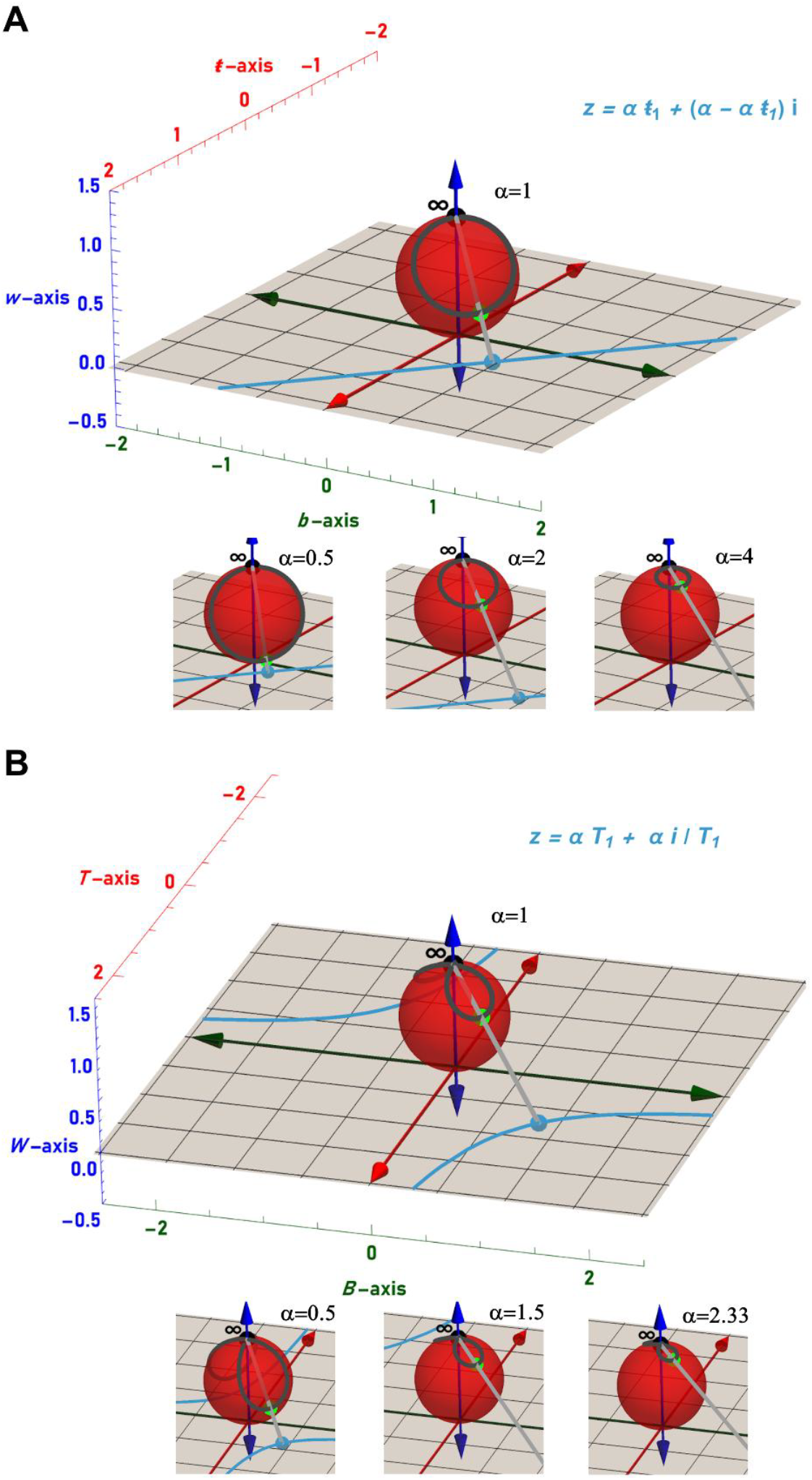
A geometric representation of the phenotypic effects of genetic modifications and/or environmental exposures on the extended complex plane (i.e., the Riemann sphere). (**A**) Stereographic projections of the reference line, Re(*z*)+Im(*z*) =1, onto a Riemann sphere (red) whose south pole is located at (0,0,0) and whose north pole is located at (0,0,1). When projected onto the surface of the sphere, the reference line forms a circle whose extent is determined by the value of the scaling factor, *a*, (compare the top panel to the three lower panels). As *a* approaches zero, the extent of the circle reaches a limit where the projection spans the full diameter of the sphere and runs through both the north and south poles. As *a* approaches infinity, the circle shrinks to become single point at the north pole. Using this representation, changes in genotype can viewed as the movement of a point along the perimeter of the circle, while changes in environment can be represented by dilations of the circle. (**B**) Stereographic projections of the reference hyperbola, Re(*Z*) ·Im(*Z*) =1, onto a Riemann sphere whose south pole is located at (0,0,0) and whose north pole is located at (0,0,1). When projected onto the surface of the sphere, the reference line forms a figure-eight (more precisely described as a lemniscate) whose extent is determined by the value of the scaling factor, *a*, (compare the top panel to the three lower panels). As *a* approaches zero the extent of the figure-eight reaches a limit where the individual loops of the projection enlarge to the point where each loop tip reaches the south pole. As *a* approaches infinity, the figure-eight shrinks to become single point at the north pole. Using this representation, changes in genotype can viewed as the movement of a point along the perimeter of the figure-eight, while changes in environment can be represented by dilations of the figure-eight.

In addition to projecting individual complex numbers, it is also possible to project the entire reference line (or reference hyperbola) to the Riemann sphere. Such projections reveal that the reference line forms a circle on the surface of the sphere, whereas the reference hyperbola takes the form of a “figure-eight” (formally referred to as a “lemniscate”) (**Figure 6A,B**). Furthermore, it is possible to reveal that changes in the value of the scaling factor, *a*, express themselves as dilations (i.e., the reduction or enlargement) of the circle or figure-eight upon the sphere surface. Decreases in *a* shift the line *z=aŧ*_*1*_+(*a*-*aŧ*_*1*_)*i*, or hyperbola *Z=aT*_*1*_*+*(*a/T*_*1*_)*i*, closer to the origin, resulting in the enlargement of the projected circle, or figure-eight. As *a* approaches zero, the circle reaches a limit where the projection spans the full diameter of the sphere and runs through both the north and south poles. On the other hand, the figure-eight reaches a limit where the individual loops of the projection enlarge to the point where each loop tip reaches the south pole. Conversely, increases in the value of *a* shift the line *z=aŧ*_*1*_+(*a*-*aŧ*_*1*_)*i*, or hyperbola *Z=aT*_*1*_*+*(*a/T*_*1*_)*i*, away from the origin and closer to infinity, resulting in the shrinking of the respective projections. Significantly, since any projected point approaches the north pole as the magnitude of *z* approaches infinity, both the circle and figure-eight shrink to a single point with coordinates (0,0,1) as *a* reaches its upper limit. This point at infinity is often referred to as an “ideal” point since its addition acts to compactify the complex plane into a finite spherical surface (**Figure 6A,B**).

As just described, the current framework thus culminates in the creation of a Riemann sphere that serves as a geometric representation of the state space of a given measurable phenotype (in relation to the phenotype exhibited by a reference genotype in a reference environment). In this way, genotypic changes can be represented geometrically as the movement of a point along either the projected circle or projected figure-eight, whereas changes in environment can be represented by dilations of the projected circle or the projected figure-eight. Remarkably, since all relevant biological factors and phenomena have precise geometric equivalents, it is possible to use such representations to unambiguously describe and visualize the effects of combinatorial changes in genetic and/or environmental factors, as well as GxE interactions. Thus, the current methodology provides a fully mature and developed tool set for genetic analysis that is both mathematically rigorous and visually intuitive. The ramifications and significance of this work within the context of the discipline of genetics, as well as with respect to the concept of measurement itself, are further explored in the Discussion below.

## Discussion

The ability to measure – i.e., the ability to describe and communicate the attributes of natural phenomena through assigning constrained numerical values to associated objects or processes – is essential to the scientific method. In its absence, there is no mechanism by which to translate the natural world into unambiguous, comparable, and communicable descriptions and thus no simple way to uncover patterns, define rules, test hypotheses, or ultimately develop comprehensive theories. The assigned numerical values are said to be constrained since they are determined through the application of distinct rule sets as described by Stevens [24]. These rules vary depending on the “resolution” needed and manifest themselves as the scale of measurement that emerges from their application.

The simplest class of scale, referred to as the “nominal” class, resolves the equality of values and is thus only capable of categorization (**Table 2**). A more sophisticated class of scale, referred to as the “ordinal” class, resolves whether a given value is greater than, or lesser than another, allowing it to rank, in addition to categorize. However, the size of the intervals between values in these scales are not known, making it impossible to determine the equality of differences. For this attribute, one must utilize the “interval” class of scales, which does indeed exhibit equal intervals and can thus be used to resolve such differences. Interval scales, though, lack a “true” zero in the sense that the zero value that is used acts only as an arbitrary reference point and does not indicate the complete absence of the thing being measured. In contrast, the most sophisticated class of scale, referred to as the “ratio” class, manifests a true or “absolute” zero and is thus capable of resolving not only the equality of differences, but also the equality of ratios. For example, when measuring temperatures using the Celsius scale (an example of an interval scale) one can correctly conclude that the interval between the values 0°C and 20°C is the same as the interval between the values 20°C and 40°C. However, one cannot conlcude that 40°C is twice as “hot” as 20°C, since 0°C does not correspond to the complete absence of thermal kinetic energy. In contrast, when using the Kelvin scale (an example of a ratio scale) one can correctly conclude that the interval between 0K and 20 K is equal to the interval between 20 K and 40K <u>and</u> that 40K is twice as hot as 20K (since 0K in this case does indeed correspond to the complete absence of thermal kinetic energy).

**Table 2:** A summary of the properties of measurement scales.

| Scale | Basic Empirical Operations | Mathematical Group Structure | Permissible Statistics (invariantive) | Other Mathematical Concepts Manifested Within the Scale | Number System |
| --- | --- | --- | --- | --- | --- |
| NOMINAL | Determination of equality | *Permutation group*<br>$x'=f(x)$<br>$f(x)$ means any one-to-one substitution | Number of cases<br>Mode<br>Contingency correlation | | Natural |
| ORDINAL | Determination of greater or less | *Isotonic group*<br>$x'=f(x)$<br>$f(x)$ means any monotonic increasing function | Median<br>Percentile | Zeroth | Whole |
| INTERVAL | Determination of equality of intervals or differences | *General linear group*<br>$x'=ax + b$ | Mean<br>Standard deviation<br>Rank-order correlation<br>Product-moment correlation | Zero (arbitrary)<br>Infinity | Real |
| RATIO | Determination of equality of ratios | *Similarity group -- SIM(1)*<br>$x'=ax$ | Coefficient of variation | Absolute Zero | Real (non-negative) |
| RELATIONAL | Determination of equality of phase | *Similarity group -- SIM(2)*<br>$z'=az$<br>where $z'$ and $z$ are complex numbers and $a$ is real | Complex mean<br>Complex standard deviation | Phase<br>Ideal Point (point at infinity) | Complex |

Importantly, each of the scales described above can be associated with a distinct number system: nominal scales with the natural numbers (where the numbers are used simply as a label or identifier), ordinal scales with the whole numbers (where the numbers are ordered, thereby permitting ranking), interval scales with the real numbers (where zero is arbitrarily chosen), and ratio scales with the non-negative real numbers (since such scales possess a true or absolute zero). Furthermore, as one progresses successively through the four classes of scales, it is possible to associate other important mathematical concepts as they appear within each class: the concept of equality in nominal scales, the concept of a zeroth value in ordinal scales (i.e., a value before the first in a series), the concept of infinity in interval scales, and finally the concept of absolute zero in ratio scales (**Table 2**). Importantly, the various scales described above make no mention of the complex numbers and thus no scale capable of determining equality of phase or manifesting the concept of an ideal point has been formally defined to date.

### Defining a “relational” scale of measurement

When viewed through the lens provided by Stevens (summarized in the previous section), the mathematical framework developed during the course of this work is easily recognizable as a formal description of a previously undefined class of measurement scale. Moreover, through the logical extension of Stevens’ principles, one can place this novel class of scale – hereafter referred to as the “relational” class – above (i.e., superseding) the ratio scale. This is because the relational class of scales manifests all the properties of the ratio class together with the capacity to resolve equality of phase (**Table 2**). This capacity, of course, cannot be achieved by any of the other scales since the notion of phase has no meaning in the context of the natural, whole, or real numbers. This is a crucial point since it is the incorporation of the concept of phase that permits the unambiguous quantification of GxG and GxE interactions (**Figures 4**,**5**). In addition to phase, the relational scale also incorporates the concept of an “ideal” point at infinity, thereby permitting division by zero and thus the ability to plot both perfect buffering or perfect transfer on the surface of the Riemann sphere (**Figure 6**).

With respect to mathematical group structure, the relational class of scales corresponds to the “similarity” group (as does the ratio class of scales). However, since ratio scales are one dimensional, they can be more precisely described as corresponding to the SIM(1) group. Likewise, since relational scales are two-dimensional, they can be more precisely described as corresponding to the SIM(2) group [25,26] (**Table 2**). Group structure, when considered in the context of measurement scales, essentially defines how a class of scale is allowed to transform its values without losing meaning. Nominal scales are the least restrictive and retain their meaning in the face of any one-to-one substitution of values. Ordinal scales, on the other hand, retain their meaning in the face of any transformation that preserves the original order of values. Interval scales retain their meaning in the face of any transformation that shifts or scales (i.e., transformations defined by *x*’=*ax*+*b*). In contrast, ratio scales retain their meaning only in the face of transformations that scale in one dimension (i.e., transformations defined by *x*’=*ax*). Finally, relational scales retain their meaning only in the face of transformations that scale in two dimensions (i.e. transformations defined by *z*’=*az*, where *z*’ and *z* are complex, and *a* is real). Thus, when applying the standards developed by Stevens, the relational scale developed during the course of this work naturally assumes a place above the ratio scale and does so through the logical extension of explicitly defined mathematical concepts (summarized in **Table 2**).

### Applying the relational scale to genetic analysis

Although many have called for drastic shifts in paradigm [1,5,27,28], it may be the case that only simple refinements to established dogma are needed to better delineate the relationship between genotypic and phenotypic variation. For example, while the relevance and importance of GxG and GxE interactions in influencing the genotype-phenotype relationship are not in question, current methods for defining the existence and magnitude of such interactions are clearly inadequate. This is primarily due to the difficulties that arise when attempting to define the “neutral” case, together with the lack of a mathematical framework that can be used to analyze the effects of both genetic variation and environmental exposures simultaneously.

The framework proposed herein provides a solution to these problem by establishing a broadly applicable method – based on the use of complex numbers – that can be used to define the expected phenotype of any GxE interaction with respect to the state space defined by the framework’s buffering parameters. Essentially, by tracking the phenotype, *x*, with respect to <u>both</u> the real and imaginary dimensions, it is possible to conceptualize a single “slice” of the phenotypic state space (defined by a given value of the scaling factor, *a*). By so doing, it is possible to unambiguously determine where on the slice *z*_obs_ falls in relation to *z*_exp_ (based on the quantity of *x* that has shifted unexpectedly between the transfer and buffer compartments due to the effects of interaction). Since such shifts are determined from the perspective of a reference genotype in a reference environment, the scale that emerges from the application of the methodology is referred to as “relational”.

This conceptualization also reveals why difficulties in defining and interpreting GxE interactions were experienced in the past. Fundamentally, such interpretations are futile when considered in the context of a one-dimensional scale based on the real numbers (i.e., a ratio scale). This is because it is only when GxE interactions are considered in the context of a two-dimensional relational scale that it becomes possible to define an <u>area</u> that can be used to quantify the interaction effects (**Eqs. 34, 37** and **Figures 4, 5**).

It should also be noted that within this conceptualization, GxG interactions are viewed simply as a particular case in which *a*=1 (i.e., when *a* is equal to one, the area quantifying an interaction effect is determined entirely by the phase difference as opposed to both a phase difference and a change in the environmental scaling factor, *a*). Thus, for the first time, it is possible to conceptualize both GxG and GxE interactions using a harmonized mathematical framework that also provides an intuitive geometric means with which to visualize interaction effects (i.e., projections on the Riemann sphere) (**Figure 6**). In any event, it is hoped that this methodology will be helpful in providing the rigorous mathematical foundation necessary to unravel the mechanisms by which GxGxE interactions influence the genotype-phenotype relationship.

## Methods

### Mathematical Derivations

#### Derivation of *A*_GxE_

As described by Zwicker [23], the signed area of a triangle on the Argand plane with vertices *z*_1_, *z*_2_, and *z*_3_ can be expressed as

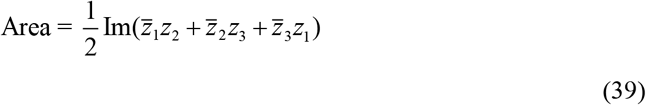

By letting *z*_1_, *z*_2_, and *z*_3_, represent the complex origin, *z*_exp_, and *z*_obs_, respectively, the area equation simplifies to Eq. (34):

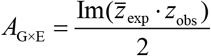

#### Derivation of *L*_disp_

The general formula for the arc length of a smooth curve from *x*_1_ to *x*_2_ is given by

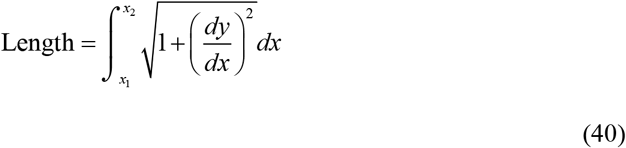

Since the derivative of *y*=*a*^2^/*x* is given by –(*a*^2^/*x*^2^) the length of the arc can be described as

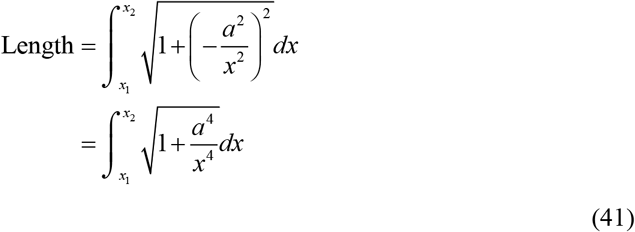

Thus, the signed length of the arc (i.e., *L*_disp_) with respect to the real axis, *T*, is given by Eq. (36):

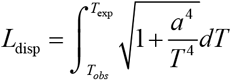

#### Derivation of *Λ*_GxE_

The definite integral of *y*=*a*^2^/*x* with respect to *x* can be expressed as

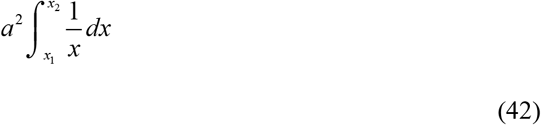

Since the definite integral of 1/*x* is ln(*x*_2_)-ln(*x*_1_), this simplifies to

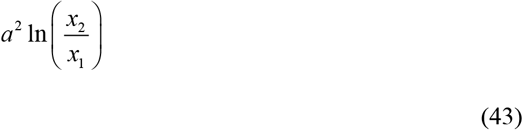

Furthermore, since the area under the curve from *x*_1_ to *x*_2_ is equivalent to the area of the region defined by the origin, *x*_1_, and *x*_2_ [29], it is thus possible to define the signed area of the hyperbolic sector as

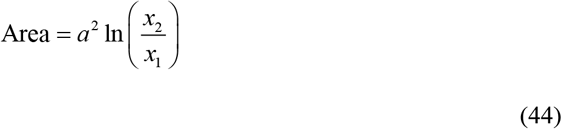

Thus, the signed area of the hyperbolic sector defined by the origin, *Z*_obs_, and *Z*_exp_ with respect to the real axis, *T* is given by Eq. (37):

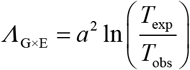

### Computational Tools

In GxE interaction plots, *A*_GxE_ and *Λ*_GxE_ are represented using heatmaps where the color gradient is determined using the “Rescale” function of the Wolfram Language (Wolfram Research Inc., Mathematica, Version 14.3. Champaign, IL). The Rescale function (https://reference.wolfram.com/language/ref/Rescale.html) outputs the provided areas rescaled from 0 to 1 over the range defined by the minimum and maximum area values. *L*_disp_ was calculated where necessary using Wolfram’s “NIntegrate” function (https://reference.wolfram.com/language/ref/NIntegrate.html).

## Funding

This work was supported by the Natural Sciences and Engineering Research Council (https://www.nserc-crsng.gc.ca/index_eng.asp) through grant R4029A03 to JK. The funders had no role in study design, data collection and analysis, decision to publish, or preparation of the manuscript.

## Competing interests

The authors have declared that no competing interests exist.

## Author contributions

Conceptualization, JK; Data curation, JK; Formal analysis, JK; Funding acquisition, JK; Investigation, JK; Methodology, JK; Visualization, JK; Writing – original draft, JK; Writing – review and editing, JK.

## Notes

### Competing Interest Statement

The authors have declared no competing interest.

## References

1. Barton NH. The “New Synthesis.” Proc Natl Acad Sci USA. 2022;119: e2122147119. doi:10.1073/pnas.2122147119

2. Benfey PN, Mitchell-Olds T. From Genotype to Phenotype: Systems Biology Meets Natural Variation. Science. 2008;320: 495–497. doi:10.1126/science.1153716

3. Gjuvsland AB, Vik JO, Beard DA, Hunter PJ, Omholt SW. Bridging the genotype– phenotype gap: what does it take? The Journal of Physiology. 2013;591: 2055–2066. doi:10.1113/jphysiol.2012.248864

4. Herrera-Luis E, Benke K, Volk H, Ladd-Acosta C, Wojcik GL. Gene–environment interactions in human health. Nat Rev Genet. 2024;25: 768–784. doi:10.1038/s41576-024-00731-z

5. Nelson RM, Pettersson ME, Carlborg Ö. A century after Fisher: time for a new paradigm in quantitative genetics. Trends in Genetics. 2013;29: 669–676. doi:10.1016/j.tig.2013.09.006

6. Orgogozo V, Morizot B, Martin A. The differential view of genotype-phenotype relationships. Front Genet. 2015;6. doi:10.3389/fgene.2015.00179

7. Baryshnikova A, Costanzo M, Kim Y, Ding H, Koh J, Toufighi K, et al. Quantitative analysis of fitness and genetic interactions in yeast on a genome scale. Nat Methods. 2010;7: 1017–1024. doi:10.1038/nmeth.1534

8. Costanzo M, Kuzmin E, Van Leeuwen J, Mair B, Moffat J, Boone C, et al. Global Genetic Networks and the Genotype-to-Phenotype Relationship. Cell. 2019;177: 85–100. doi:10.1016/j.cell.2019.01.033

9. Mani R, St.Onge RP, Hartman JL, Giaever G, Roth FP. Defining genetic interaction. Proc Natl Acad Sci USA. 2008;105: 3461–3466. doi:10.1073/pnas.0712255105

10. Waddington CH. The Strategy of the Genes. 0 ed. Routledge; 2014. doi:10.4324/9781315765471

11. Manuck SB. The reaction norm in gene × environment interaction. Mol Psychiatry. 2010;15: 881–882. doi:10.1038/mp.2009.139

12. Moore DS. Gene × Environment interaction: What exactly are we talking about? Research in Developmental Disabilities. 2018;82: 3–9. doi:10.1016/j.ridd.2018.04.012

13. Ottman R. Gene-environment interaction: definitions and study designs. Prev Med. 1996;25: 764–770. doi:10.1006/pmed.1996.0117

14. Tabery J. R. A. Fisher, Lancelot Hogben, and the Origin(s) of Genotype–Environment Interaction. J Hist Biol. 2008;41: 717–761. doi:10.1007/s10739-008-9155-y

15. Karagiannis J. A mathematical framework for the quantitative analysis of genetic buffering. Cordell HJ, editor. PLoS Genet. 2025;21: e1011730. doi:10.1371/journal.pgen.1011730

16. Schmitt BM. The quantitation of buffering action I. A formal & general approach. Theor Biol Med Model. 2005;2: 8. doi:10.1186/1742-4682-2-8

17. Schmitt BM. The quantitation of buffering action II. Applications of the formal & general approach. Theor Biol Med Model. 2005;2: 9. doi:10.1186/1742-4682-2-9

18. Baryshnikova A, Costanzo M, Myers CL, Andrews B, Boone C. Genetic Interaction Networks: Toward an Understanding of Heritability. Annu Rev Genom Hum Genet. 2013;14: 111–133. doi:10.1146/annurev-genom-082509-141730

19. Bak J, Newman DJ. Complex analysis. Third edition. New York Dordrecht Heidelberg London: Springer; 2017.

20. Mathews JH, Howell RW. Complex analysis for mathematics and engineering. 3. ed. Boston: Jones and Bartlett; 1997.

21. Needham T. Visual complex analysis. Oxford: Clarendon Press; 1997.

22. Ponce Campuzano, J. C. Complex Analysis: A Visual and Interactive Introduction. 2019. Available: https://complex-analysis.com/

23. Zwikker C. The advanced geometry of plane curves and their applications. Mineola, NY: Dover Publications; 1963.

24. Stevens SS. On the Theory of Scales of Measurement. Science. 1946;103: 677–680.

25. Hawkins T. The Erlanger Programm of Felix Klein: Reflections on its place in the history of mathematics. Historia Mathematica. 1984;11: 442–470. doi:10.1016/0315-0860(84)90028-4

26. Rowe DE. Felix Klein: The Erlangen Program. Cham: Springer Nature Switzerland; 2025. doi:10.1007/978-3-031-85474-3

27. Tautz D, Pallares LF, Andersson L, Barghi N, Barton N, Bay R, et al. Beyond Mendel: a call to revisit the genotype–phenotype map through new experimental paradigms. Long A, editor. GENETICS. 2026;232: iyag024. doi:10.1093/genetics/iyag024

28. Huang S, Soto AM, Sonnenschein C. The end of the genetic paradigm of cancer. PLoS Biol. 2025;23: e3003052. doi:10.1371/journal.pbio.3003052

29. Nelsen RB. The Relationship Between Hyperbolic and Exponential Functions. The College Mathematics Journal. 1988;19: 54–56. doi:10.1080/07468342.1988.11973091

